# Striking differences in virulence, transmission, and sporocyst growth dynamics between two schistosome populations

**DOI:** 10.1101/685958

**Authors:** Winka Le Clec’h, Robbie Diaz, Frédéric D. Chevalier, Marina McDew-White, Timothy J.C. Anderson

## Abstract

**Background:** Parasite traits associated with transmission success, such as the number of infective stages released from the host, are expected to be optimized by natural selection. However, in the trematode parasite *Schistosoma mansoni*, a key transmission trait – the number of cercariae larvae shed from infected *Biomphalaria spp*. snails – varies significantly within and between different parasite populations and selection experiments demonstrate that this variation has a strong genetic basis. In this study, we compared the transmission strategies of two laboratory schistosome population and their consequences for their snail host.

**Methods:** We infected inbred *Biomphalaria glabrata* snails using two *Schistosoma mansoni* parasite populations (SmBRE and SmLE), both isolated from Brazil and maintained in the laboratory for decades. We compared life history traits of these two parasite populations by quantifying sporocyst growth within infected snails (assayed using qPCR), output of cercaria larvae, and impact on snail host physiological response (i.e. hemoglobin rate, laccase-like activity) and survival.

**Results:** We identified striking differences in virulence and transmission between the two studied parasite populations. SmBRE (low shedder (LS) parasite population) sheds very low numbers of cercariae, and causes minimal impact on the snail physiological response (i.e. laccase-like activity, hemoglobin rate and snail survival). In contrast, SmLE (high shedder (HS) parasite population) sheds 8-fold more cercariae (mean ± se cercariae per shedding: 284±19 vs 2352±113), causes high snail mortality, and has strong impact on snail physiology. We found that HS sporocysts grow more rapidly inside the snail host, comprising up to 60% of cells within infected snails, compared to LS sporocysts which comprised up to 31%. Cercarial production is strongly correlated to the number of *S. mansoni* sporocyst cells present within the snail host tissue, although the proportion of sporocyst cells alone does not explain the low cercarial shedding of SmBRE.

**Conclusions:** We demonstrated the existence of alternative transmission strategies in the *S. mansoni* parasite consistent with trade-offs between parasite transmission and host survival: a “boom-bust” strategy characterized by high virulence, high transmission and short duration infections and a “slow and steady” strategy with low virulence, low transmission but long duration of snail host infections.

## BACKGROUND

Models predicting the evolution of virulence for parasites transmitted horizontally assume generally that transmission rate (i.e. the probability for an infected host to infect a susceptible new host) and virulence (i.e. the increase in host mortality due to infection) are positively correlated, as higher production of infective stages may be more harmful for the host [1–4]. For most of the virulence evolution models, such a trade-off shapes the relationship between parasite transmission and host survival (the higher the virulence of the parasite, the shorter the host survival, and in turn the parasite lifespan) [5] for both micro [6] and macroparasites [7]. This transmission / virulence trade-off model provides a general and intuitive framework for understanding within-species variation in parasite virulence [5,8].

Studying production of larvae of schistosome, a human trematode parasite, offers the dual benefits of empirically testing the trade-off model for a macroparasite and improving our understanding of a key transmission-related trait in a biomedically important helminth parasite. Schistosomes infect over 200 million people in 78 countries (WHO fact sheet No. 115, http://www.who.int/mediacentre/factsheets/fs115/en/), causing schistosomiasis. This chronic and debilitating tropical disease ranks second behind malaria in terms of morbidity and mortality; there is no licensed vaccine and only one drug (Praziquantel) is available to treat patients. Schistosome parasites have a complex lifecycle, involving a freshwater snail (intermediate host) and a mammal (definitive host). When parasite eggs are expelled with mammal feces or urine in water, miracidia larvae hatch and actively search for its snail vector. Larvae penetrate the snail head-foot, differentiate into mother sporocysts and then asexually proliferate to generate daughter sporocysts. This intramolluscan parasite stage, while growing, metabolizes snail tissues, such as the hepatopancreas and the albumen gland [9]. These organs are involved in the protein and egg production, and schistosome infection results in castration of infected snails [10]. After approximately a month of infection, daughter sporocysts start to release cercariae, the mammal infective larval stage of the parasite. These exit through the snail body wall and are released into the water. This complex lifecycle can be maintained in the laboratory using rodent definitive hosts and freshwater snail intermediate hosts.

Lewis and colleagues [11] measured production of *S. mansoni* cercariae from infected *Biomphalaria glabrata* snails in the laboratory, and determined that this transmission trait varies significantly within and between different parasite populations. Moreover, Gower and Webster [12] performed replicated selection experiments in the laboratory and showed that cercarial shedding from the snail host responded extremely rapidly to selection, with a 7-fold change in cercarial production within three generations. These observations suggest that variation in transmission stage production in *S. mansoni* has a strong genetic basis. Following the transmission / virulence trade-off model, we hypothesized that *S. mansoni* parasites producing many cercariae will negatively affect snail health and cause high virulence. Virulence could occur because intramolluscan schistosome stages consume host tissue to produce large numbers of cercariae, and/or because cercariae damage tissue when they are released from snails. On the other hand, parasites that produce less larvae, but for a longer period of time, will be less virulent toward their snail host and have a lower negative impact on their physiology and survival.

In this study, we investigated the transmission and virulence of two laboratory *S. mansoni* populations both originating from South America. We observed striking differences in the number of cercariae produced by these two populations of schistosome parasites and showed that these transmission-related life-history traits have a genetic basis. We then investigated why cercarial production varies between these two populations by investigating growth of sporocysts within each infected snails. Finally, we highlight a negative relationship between transmission stage production and snail survival, health and immune parameters. Our results support the presence of a virulence / transmission trade-off in *S. mansoni* / *B. glabrata*.

## METHODS

### Ethics statement

This study was performed in strict accordance with the recommendations in the Guide for the Care and Use of Laboratory Animals of the National Institutes of Health. The protocol was approved by the Institutional Animal Care and Use Committee of Texas Biomedical Research Institute (permit number: 1419-MA-3).

### Biomphalaria glabrata s*nails and* Schistosoma mansoni *parasites*

Uninfected inbred albino *Biomphalaria glabrata* snails (line Bg26 derived from 13-16-R1 line [13] were reared in 10-gallon aquaria containing aerated freshwater at 26-28°C on a 12L-12D photocycle and fed *ad libitum* on green leaf lettuce. All snails used in this study had a shell diameter between 8 and 10 mm, as snail size can influence cercarial outcome [14,15]. For all the experiments presented in this study, we used inbred snails to minimize the impact of snail host genetic background on the parasite life history traits. *B. glabrata* snails Bg26 were inbred over 3 generations through selfing [13], so their genomes are expected to be 87.5% identical by descent.

The SmLE schistosome population (high shedder, HS) was originally obtained by Dr. J. Pellegrino from an infected patient in Belo Horizonte (Minas Gerais, Brazil) in 1965 and has since been maintained in laboratory [11], using *B. glabrata* NMRI population as intermediate host and Syrian golden hamster (*Mesocricetus auratus*) as definitive hosts. The SmBRE schistosome population (low shedder, LS) was sampled in the field in 1975 from Recife (East Brazil) [16] and has been maintained in the laboratory in its sympatric albino Brazilian snail host BgBRE using hamsters or mice as the definitive host.

### *Measurement of* S. mansoni *life history traits and virulence*

We compared larval output (i.e. cercarial production) and intramolluscan development (i.e. sporocyst development and growth) of SmLE (HS) and SmBRE (LS) parasite populations using the same inbred snail population in two independent cohort experiments (Figure 1). We also measured the impact of these parasitic infections on the snail host by quantifying snail survival and physiological responses (i.e. laccase-like activity and hemoglobin rate in the hemolymph).

**Figure 1:**
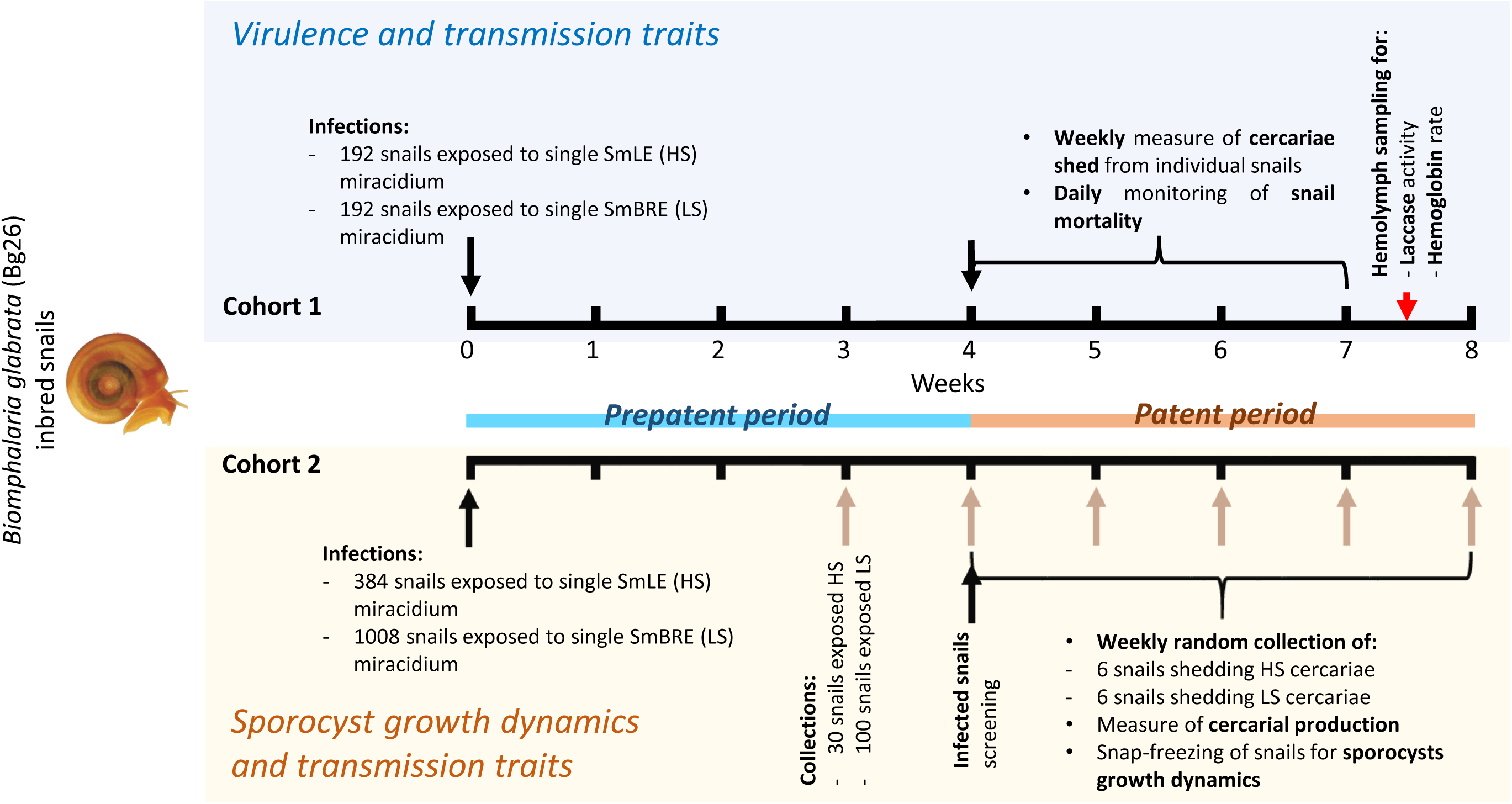
Outline of the experimental design. We used two independent cohorts of *Biomphalaria glabrata* Bg26 inbred snails. Each snail was exposed to one miracidium from the SmLE (HS) or SmBRE (LS) *Schistosoma mansoni* populations. In cohort 1, we measured transmission stage production for SmLE (HS) and SmBRE (LS) populations during 4 weeks of the patent period (week 4 to 7 post-infection). We also evaluated the virulence of these two populations of parasite by measuring the daily snail survival during the patent period. After 7.5 weeks post-infection, surviving infected snails were bled and we measured the total laccase-like activity as well as the hemoglobin rate in the collected hemolymph samples. We used cohort 2 to determine the weekly sporocyst growth dynamics in snails for the late prepatent (week 3) and the patent period (week 4 to 8).

#### 1. Cohort 1: S. mansoni *cercarial output and impact on snail survival and physiological response*

##### a. S. mansoni cercarial production over time

To compare cercarial production over the time for the SmLE (HS) and SmBRE (LS) parasite populations, we exposed 384 inbred *B. glabrata* Bg26 to single miracidia. Miracidia of each population were hatched from eggs recovered from 45-day-infected female hamster livers infected. The livers were homogenized and the eggs were filtered, washed with normal saline (154 mM sodium chloride (Sigma), pH 7.5), transferred to a beaker containing freshwater, and exposed to artificial light to induce hatching. We exposed individual snails (192 per parasite populations) to single miracidia in 24 well-plates for at least 4 hours, and exposed snails were maintained in trays (48 per tray) for 4 weeks. We used single miracidia for infections to avoid competition effects and obtain the phenotype corresponding to a single parasite genotypes. We covered trays with black plexiglass lids after 3 weeks to reduce cercarial shedding. At four weeks post-exposure, we placed snails in 1 mL freshwater in 24 well-plates under artificial light for 2 hours to induce cercarial shedding. We isolated each infected snail in a 100 mL glass beaker filled with ∼50 mL freshwater and kept them in the dark until week 7 post-infection. We replaced freshwater as needed (typically every two days) and fed snails *ad libitum*. To quantify cercarial shedding, we placed infected snails in 1 mL freshwater in a 24 well plate under artificial light (as described above) every week for 4 weeks (week 4 to 7). For each well, we sampled three 100 µL aliquots, added 20 µL of 20X normal saline and counted the immobilized cercariae under a microscope. We multiplied the mean of the triplicated measurement by the dilution factor (10) to determine the number of cercariae produced by each infected snail. We also extracted DNA from week 4 cercariae to determine parasite gender by PCR [17].

##### b. B. glabrata snail survival

We evaluated survival of the infected snails in cohort 1 over the course of the infection. We compared the survival of infected snails with a group of 48 uninfected control snails. We monitored snail survival every day from the first cercarial shedding day (28 days after exposure) to 22 days after the first cercarial shedding (50 days after exposure).

##### c. Snail physiological response to parasitic infection: hemolymph laccase-like activity and hemoglobin rate

In week 7, 3 days after the last cercarial shedding, we collected hemolymph as described in Le Clec’h et al., 2016 [18] from all surviving snails infected with SmLE (HS) and SmBRE (LS), as well as from uninfected Bg26 controls maintained under the same conditions. We measured both laccase-like activity (phenoloxydase (PO) activity, known to be involved in invertebrate immunity) and the hemoglobin (protein carrying oxygen in *B. glabrata* hemolymph) rates in the hemolymph of each snail infected with SmLE (HS), SmBRE (LS) or uninfected control.

We measured the total laccase-like activity as described in Le Clec’h et al., 2016 [18]. In brief, we combined 10 µL of freshly collected hemolymph to 40 µL of cacodylate buffer and 40 µL of bovine trypsin (1mg/mL) in a 96-well optical plate (Corning). Each sample was coupled to a control where 10 µL of the same hemolymph aliquot was combined to 40 µL of 10 mM diethylthiocarbamate (a specific inhibitor of PO enzymes) and 40 µL of bovine trypsin. We incubated plates in the dark, at 37°C for 45 minutes, then added 120 µL of the freshly prepared *p*-phenylenediamine substrate (50 mM). After 2 hours incubation at 37°C, before reaching the plateau phase of the laccase-like activity, we measured dopachrome formation at λ=465 nm using a SpectraMax M1 (Molecular Devices). A substrate auto-oxidation control was also performed for each experiment, where the hemolymph sample was replaced by 10 µL of distilled water. The value of this substrate auto-oxidation control were subtracted from sample and control values.

To quantify the hemoglobin rate, we centrifuged the hemolymph (5 minutes, at 300 × g at 4°C) to pellet the hemocytes (i.e. the immune cells), as the hemoglobin is not sequestrated in cells but free in the plasma [19]. We collected the plasma in a fresh microtube placed on ice. The hemoglobin rate was determined by measuring the optical density (OD) of the hemoglobin solution at λ=410 nm, the maximum absorption of *B. glabrata* hemoglobin [19]. In a 96-well optical plate (Corning), we combined 190 µL of PBS 1X to 10 µL of plasma. A blank control, containing only 200 µL of PBS 1X was also performed for each assays. The value obtained for this blank control was subtracted from the sample wells (i.e. containing plasma).

#### 2. Cohort 2: S. mansoni *sporocyst time course and growth dynamics*

##### a. Cohort design

Evaluating sporocyst development within infected snails requires the sacrifice of snails at multiple time points. We exposed a second cohort of 1,392 Bg26 inbred snails to single miracidia from our two *S. mansoni* populations. We exposed 384 snails to SmLE (HS) and 1,008 to SmBRE (LS). The numbers are unequal because SmLE (HS) shows much higher infection rates than SmBRE (LS). Snails were kept in trays (48 per tray). We sampled snails at week 3 post-exposure, one week prior to cercarial maturation. To ensure sampling of infected snails, we randomly picked 30 snails exposed to SmLE (HS) and 100 snails exposed to SmBRE (LS). These sampling numbers take in account the snail susceptibility to 1 parasite larva.

In week 4, we placed each infected snail in 1 mL of freshwater in 24-well plates under artificial light for 2 hours to induce cercarial shedding and identify infected snails. From week 4-8, we isolated infected snails in trays (48 per tray) under a black lid. Each week, we randomly picked 6 infected snails from each parasite population and counted the cercariae released by each snail as described in section 1a.

We cleaned the shell of sampled snails with 70% ethanol, snap-froze them individually in liquid nitrogen, and store them at −80°C for further molecular analysis.

##### b. gDNA extractions from exposed snails

To prepare exposed snails for molecular analysis, we crushed snails individually in a sterile, liquid nitrogen-cooled mortar and pestle to create a fine, homogenized tissue powder, and kept 100 µL of powder into 1.5 mL tubes at −80°C until gDNA extraction. We extracted the gDNA using a DNeasy Blood & Tissue Kit (Qiagen) according to manufacturer instructions, with tissue lysis for 20 minutes at 56°C. We quantified the gDNA using a Qubit dsDNA BS Assay Kit (Invitrogen).

##### c. Multiplex PCR to identify infected snails

To screen for infected prepatent snails (week 3, Figure 1), we performed a multiplex PCR on the gDNA recovered from snails powder. We used the *α-tubulin 2 S. mansoni* gene (accession number S79195.1; gene number Smp_103140; [20]) as specific parasite marker and the P-element induced wimpy testis (*piwi*) gene from *B. glabrata* [21] as specific snail host marker. We identified the *piwi* gene (BGLTMP009852) in the *B. glabrata* genome (BglaB1 assembly) using the blast module of VectorBase [22] and 3 ESTs showing similarities with *piwi* (accession numbers FC855819.1, FC856421.1, and FC856380.1).

Multiplex PCR reactions consisted of 8.325 µL sterile water, 1.5 µL 10x buffer, 1.2 µL dNTP (2.5 mM each), 0.9 µL MgCl_2_, 0.5µL of each primer (10 µM) for both markers (*piwi* F: 5’-CTTCTCCAATGCTACCATCAAAG-3’; *piwi* R: 5’-TTTCATCCTCCACACTGACAA-3’; *α-tubulin 2* F: 5’-CGACTTAGAACCAAATGTTGTAGA-3; *α-tubulin 2* R: 5’-GTCCACTACATTGATCCGCT-3’), 0.075 µL of *Taq* polymerase (TaKaRa) and 1µL of gDNA template using the following program: 95°C for 5 minutes, [95°C for 30s, 55°C for 30s, and 72°C for 30s] × 35cycles, 72°C for 10 minutes. Infected snails exhibit a two band pattern at 361 bp and 190 bp on an agarose gel while uninfected snails show one band at 361 bp (Supplementary figure 1). All the primers were designed using PerlPrimer v1.21.1 [23].

##### d. qPCR to quantify the proportion of sporocyst cells within infected snails

The daughter sporocysts that release cercariae are intertwined in the snail tissue, making them difficult to isolate and study so they have been neglected relative to other parasite life stages. Using a custom quantitative PCR assay, we quantified the relative proportion of parasite cells within infected snail at different time points of the infection. This qPCR assay provides a relative measure of parasite growth within infected snails.

We quantified a single copy gene from the parasite (*α-tubulin 2*, see section 2.c, [20]) and from the snail (*piwi*, see section 2.c, [21]). The qPCR assay used a different set of *piwi* primers (*piwi* F [5’-AATCATCTCATTCAACCTGTCCAT-3’] and *piwi* R [5’-ATTTCCGCCATCATAGCCC-3’]) amplifying a 107 bp amplicon and the same *α-tubulin 2* primers as described in the end-point PCR assay. We conducted qPCR in duplicate for each reaction (i.e. samples and standards). Reactions consisted of 5 µL SYBR Green PCR master mix (Applied Biosystems), 3.4 µL sterile water, 0.3 µL of each primer (10 µM) and 1µL of standard PCR product or sample gDNA. We used the following program: 95°C for 10 minutes, [95°C for 15s and 60°C for 1 minute] × 40 cycles followed by a melting curve step (15s at 95°C and then rising in 0.075°C increments/second from 60°C to 95°C), to check for the uniqueness of the product amplified. We plotted standard curves using seven 10-fold dilutions of a purified α-tubulin 2 PCR product for *S. mansoni* parasite (*α-tubulin 2* copies.µL^−1^: 2.69×10^1^ - 2.69×10^7^) and seven 10-fold dilutions of a purified *piwi* PCR product for *B. glabrata* (*piwi* copies.µL^−1^: 2.60×10^1^ - 2.60×10^7^). PCR products for standard curves were generated using TaKaRa Taq R001 AM kit (Clonetech) and the manufacturer’sprotocol (PCR cycles: 95° for 5 min, [95°C for 30 s, 60°C for 30 s, 72°C for 30 s] × 35 cycles, 72° for 10 min), purified using SigmaSpin Sequencing Reaction Clean-Up kit (Sigma) following the manufacturer’s protocol, and quantified using Qubit dsDNA BR Assay kit (Invitrogen). We estimated the number of copies in the PCR products as follows: PCR product length × (average molecular mass of nucleotides (330 g.mol^−1^) × 2 strands) × Avogadro constant. The number of *α-tubulin 2* and *piwi* copies in each sample was estimated according to the standard curve (QuantStudio Design and Analysis Software). Both snail and parasite genes quantified are present as a single copy gene, so the number of gene copies quantified corresponds to the number of genomes of each organism. As both parasite and snail are diploid, the number of genomes is directly proportional to the number of cells from each organism. The proportion of parasite cells within infected snails, our relative measure of parasite growth, was calculated as follows:

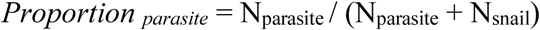

Where N is the number of parasite or snail cells measured by qPCR.

### Statistical analysis

All statistical analyzes and graphs were performed using R software (version 3.5.1). When data were not normally distributed (Shapiro test, p < 0.05), results were compared with a Kruskal-Wallis followed by Dunn’s multiple comparison test or simple pairwise comparison (Wilcoxon-Mann-Whitney test). When data followed a normal distribution, results were compared with a simple pairwise comparison Welsh *t*-test. We performed survival analysis using log-rank tests (R survival package) and correlations analysis with Pearson’s test. The confidence interval of significance was set to 95% and p-values less than 0.05 were considered as significant.

## RESULTS

### Striking differences in transmission stage production between two *Schistosoma mansoni* populations

The SmLE (HS) parasite population produce more cercariae than the SmBRE (LS) population in both cohort 1 (4 weeks of shedding) and cohort 2 (5 weeks of sheddings): on average, our SmLE (HS) population shed 8-fold more cercariae than SmBRE (LS) (Cohort 1: Kruskal-Wallis test, *p* < 2.2 × 10^−16^, Figure 2A; Cohort 2: Kruskal-Wallis test, *p*=2.824 × 10^−11^, Figure 3B). In this experiment, all the infected snails were from the same inbred *B. glabrata* population (Bg26) to minimize the impact of the host genetic background, because we know that cercarial shedding can be influenced by the snail genotype [24].

**Figure 2:**
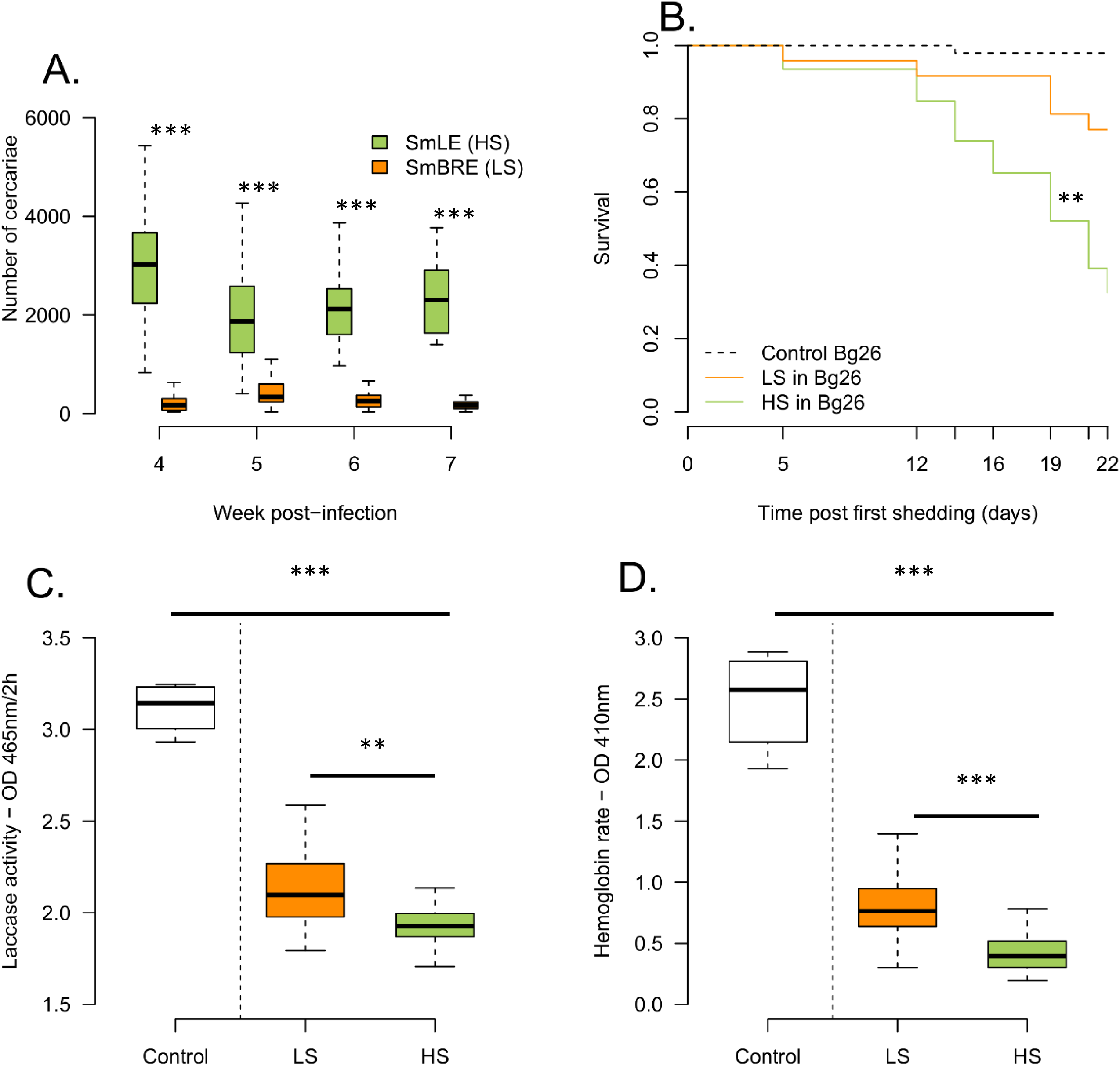
Transmission stage production and virulence of two *Schistosoma mansoni* populations (SmLE (HS) and SmBRE (LS)). **(A)** Difference in the number of cercariae produced by SmLE (HS) and SmBRE (LS) *S. mansoni* populations during 4 weeks of the patent period (week 4 to 7 post infection). SmLE (HS) population is shedding more cercariae than SmBRE (LS) population of parasite at all the time points. **(B)** Survival of the infected and control *Biomphalaria glabrata* (Bg26 inbred) snails from the first day of cercarial shedding to day 22 after the first shedding. Infection with SmLE (HS) parasites results in greater snail mortality than infection with SmBRE (LS) parasites. **(C)** Infected snails show a decrease in laccase-like activity in the snail hemolymph compared to uninfected ones. Snails infected with SmLE (HS) parasites show a greater decrease than that in snails parasitized by SmBRE (LS) parasites. **(D)** The overall hemoglobin rate in the hemolymph is reduced by the presence of schistosome parasites. However, the reduction is greater when snails are infected with the SmLE (HS) parasites. \**p* < 0.05; ** *p* ≤ 0.01; *** *p* ≤ 0.001.

**Figure 3:**
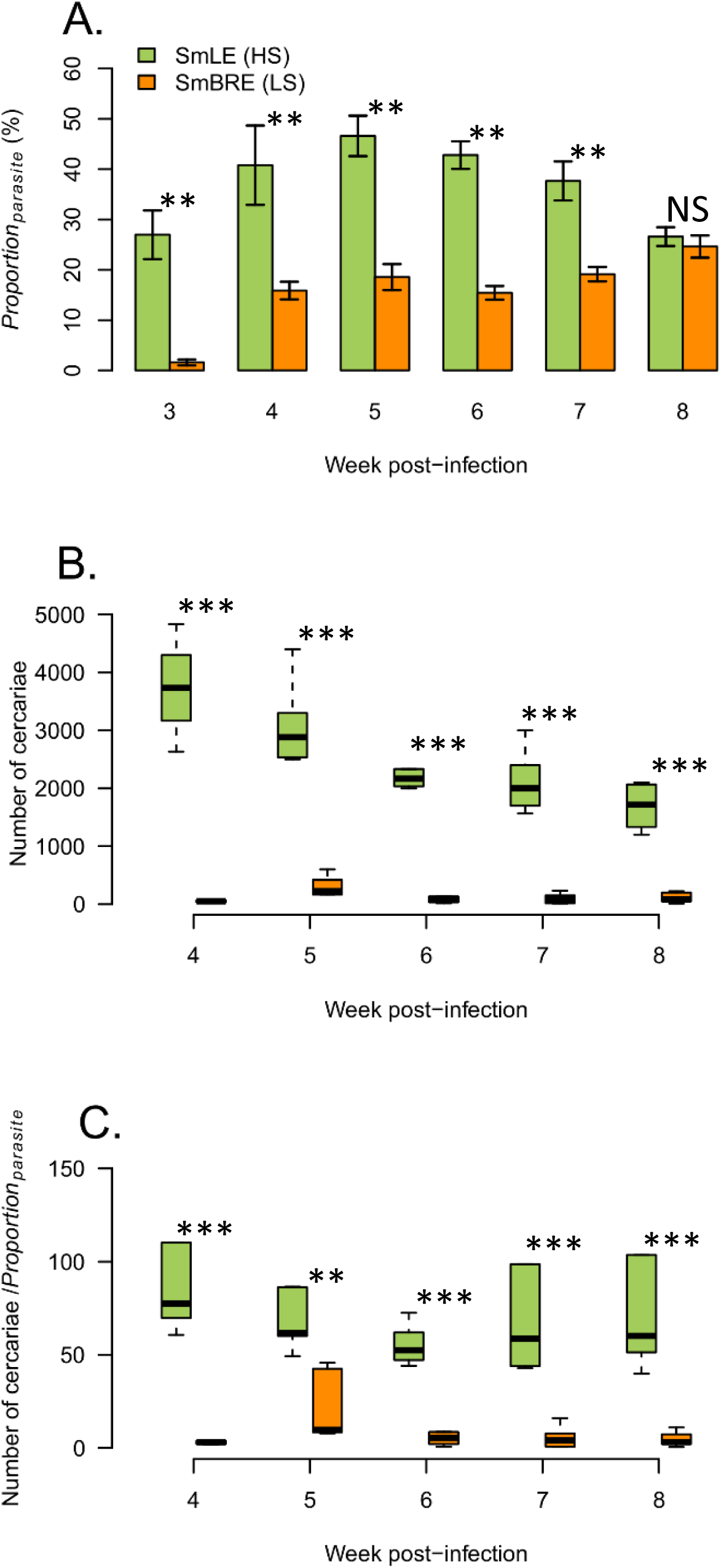
Sporocyst growth dynamics and cercarial production in SmLE (HS) and SmBRE (LS) *S. mansoni*. **(A)** Comparison of the daughter sporocyst developmental kinetics for SmLE (HS) and SmBRE (LS). The proportion of sporocyst cells within snails (*Proportion*_*parasite*_) were quantified by qPCR during 6 weeks of the infection (from week 3 to 8). SmLE (HS) sporocysts grow faster and are more numerous than the SmBRE (LS) ones. **(B)** Cercarial shedding profiles of the SmLE (HS) and SmBRE (LS) during the 5 weeks of the patent period (i.e. cercarial shedding time) (weeks 4 to 8). SmLE (HS) parasites produce significantly more cercariae than the SmBRE (LS) parasites. **(C)** Ratio calculated by dividing the number of cercariae produced by the proportion of daughter sporocyst cells present in the snail, infected with SmLE (HS) or the SmBRE (LS) population. Differences in proportions of sporocyst within infected snails are not sufficient to explain the difference in cercarial output between the SmLE (HS) and SmBRE (LS) infected snails. \**p* < 0.05; ** *p* ≤ 0.01; *** *p* ≤ 0.001.

We exposed the snail hosts to only one miracidia (SmLE (HS) or SmBRE (LS)), male or female. We were therefore able to sex each parasite that developed inside the host and test the influence of the parasite gender on the cercarial output. Parasite sex does not impact the cercarial production in the SmLE (HS) population (Wilcoxon test, *p*=0.6; Supplementary figure 2) but does have an influence on the SmBRE (LS) population where male LS genotypes produced significantly less cercariae than female ones (Wilcoxon test, *p*=0.016; Supplementary figure 2).

### *Increased virulence of the high shedder* S. mansoni *population*

#### 1. Comparison of survival rates in infected snails

SmLE (HS) parasites have a strongest impact on the survival of the host after the first shedding day (log-rank test global analysis: *p*=8 × 10^−4^) compared to SmBRE (LS) (log-rank test: *p*=0.011) or the control group (i.e. uninfected snails; log-rank test: *p*=0.004). Snails infected with SmBRE (LS) parasite did not have significantly greater mortality than controls (Figure 2B).

#### 2. Comparison of hemoglobin and laccase-like activity in infected snails

We also measured the impact of our two population of parasites on their snail hosts by measuring laccase-like activity in the snail hemolymph, 7.5 weeks after parasite exposure [18]. Unlike the survival data, where only SmLE (HS) population has a negative impact on the snail host, we observed that the laccase-like activity is reduced in snails infected with both SmBRE (LS) and SmLE (HS) relative to controls (Kruskal-Wallis test, *p*=1.108 × 10^−5^; Figure 2C). However, snails infected with SmLE (HS) population showed greater reduction in laccase-like activity than those infected with SmBRE (LS) (Welsh *t*-test, *p*=0.001; Figure 2C).

We observed a similar impact of infection on hemoglobin rate, measured in hemolymph samples collected 7.5 weeks after parasite exposure. Both SmLE (HS) and SmBRE (LS) infected snails had reduced hemoglobin relative to controls (Kruskal-Wallis test, *p*=2.155 × 10^−7^; Figure 2D), while SmLE (HS) infected snails had significantly reduced hemoglobin relative to SmBRE (LS) infected snails (Wilcoxon test, *p*=3.692 × 10^−6^; Figure 2D).

Moreover, we found a strong positive correlation between hemoglobin rate and laccase-like activity (Pearson’s correlation test: 0.78, *p*=4.633 × 10^−12^, Supplementary figure 3C). These two proteins provide a good proxy of snail health and are both severely impacted by *S. mansoni*, with the impact dependent on parasite population (SmBRE (LS) or SmLE (HS)). We observed a strong negative correlation between the hemoglobin rate and the average number of cercariae produced (Pearson’s correlation test: −0.54, *p*=2.528 × 10^−5^, Supplementary figure 3B) as well as between the laccase-like activity in the hemolymph and the average number of cercariae produced (Pearson’s correlation test: − 0.33, *p*=0.017, Supplementary figure 3A). These physiological proxies of snail health support the mortality data, showing that our SmLE (HS) population of *S. mansoni* is more virulent toward the snail inbred host (Bg26) than the SmBRE (LS) population of parasites.

### Dynamics of sporocyst growth in infected snails

Cercariae are free-living schistosome larvae produced by daughter sporocysts. This parasite stage is intertwined with the snail host hepatopancreas and ovotestis, so is difficult to quantify even in dissected snails. In cohort 2, we investigate the relationship between the quantity of cercariae released and the total quantity of sporocyst tissue developing in the snail for both SmLE (HS) and SmBRE (LS) parasites. We measured the proportion of parasite cells relative to snail cells (*Proportion*_*parasite*_) within infected snails using a custom qPCR assay. SmLE (HS) parasites have a significantly higher growth (average *Proportion*_*parasite*_ ranges from average values of 26 % to 47 % with a maximum of 60.46 % for individual snails) than SmBRE (LS) parasites (average *Proportion*_*parasite*_ ranges from average values of 1.5 % to 25 %, with a maximum of 31.12% for individual snails) across the time course experiment (Kruskal-Wallis test, *p*=1.64 × 10^−7^, Figure 3A). However, at the end of the time course (week 8 post-infection) both SmBRE (LS) and SmLE (HS) sporocysts comprise approximately 25% of snail tissue (Welsh *t*-test, *p*=0.510; Figure 3A). The sporocyst growth profiles also differ in shape: SmLE (HS) reaches a peak in week 5 during the second week of shedding and then declines, while SmBRE (LS) is still increasing in week 7 at the end of the timecourse.

Differences in *Proportion*_*parasite*_ explains some, but not all of the variation in cercarial shedding between the SmLE (HS) and SmBRE (LS) populations. At week 8, values of *Proportion*_*parasite*_ are similar for SmLE (HS) and SmBRE (LS), but SmLE (HS) still continues to produce around 15 times more cercariae than SmBRE (LS) (mean ± se cercariae: SmLE (HS): 1689±164 vs SmBRE (LS): 109±35). Furthermore, in weeks 4 to 8, the differences in *Proportion*_*parasite*_ are not sufficient to explain the differences in cercarial output between the SmLE (HS) and SmBRE (LS) infected snails (Kruskal-Wallis test, *p* = 5.22 × 10^−11^, Figure 3C).

We observed a strong correlation between *Proportion*_*parasite*_ and the quantity of cercariae released by the same snail (Pearson’s test, coef. = 0.77, *p*=1.058 × 10^−12^, Figure 4A). This correlation is driven by SmLE (HS) parasite population (Figure 4B), and there was no correlation observed for SmBRE (LS), for which there is more limited variation in the number of cercariae produced (Figure 4C).

**Figure 4:**
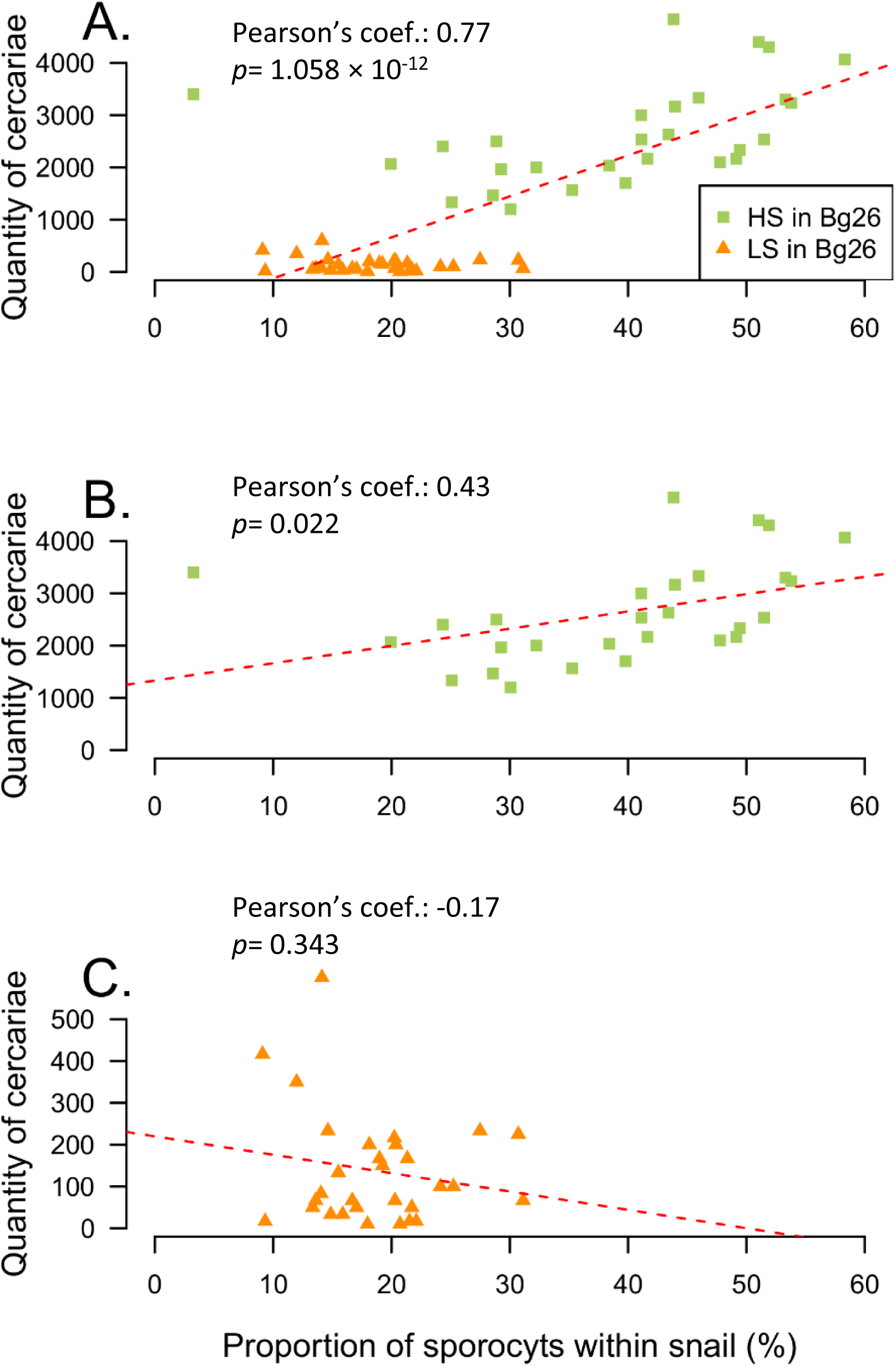
Correlations between sporocyst quantity and cercarial output for SmLE (HS) and SmBRE (LS) parasites. **(A)** There is a strong correlation between the proportion of sporocysts present in the snail tissue and the quantity of cercariae released by the same snail (Pearson’s test, coef. = 0.77). This correlation is mainly driven by **(B)** the SmLE (HS) parasite population (Pearson’s test, coef. = 0.43) **(C)** There is no significant correlation for the SmBRE (LS) parasite population (Pearson’s test, coef. = −0.17).

## DISCUSSION

### Virulence – transmission trade-offs in *S. mansoni*

Our results are consistent with a transmission / virulence trade-off model [25–29]. We show that the quantity of sporocysts, the intramolluscan stage of *S. mansoni*, and the number of cercariae shed are strongly correlated. The SmLE (HS) parasite population has a higher sporocyst growth rate within infected snails, produces large numbers of cercariae (i.e. transmission stage) and it is highly virulent toward the snail. Virulence was evident both from the high rate of snail mortality and from reduced levels of two physiological parameters: hemoglobin rate [30,31] and laccase-like activity [18]. In contrast, the SmBRE (LS) population of *S. mansoni* has a much lower sporocyst growth rate inside the snail host, releases fewer cercariae compared to the SmLE (HS) parasite, and is much less virulent for its snail host. Similar patterns of life history variation are also observed in the water flea/bacteria (*Daphnia magna* / *Pasteuria ramosa)* infection model, a comparable system where bacterial infection castrates water fleas. In this system, water fleas infected with high virulent “early killer” spores had a significantly higher death rate compared to those infected with low virulence “late killers”. Variation in time of death was at least in part caused by genetic differences among parasites [6].

Our findings provide an interesting contrast with patterns observed in experimental work on the SmPR1 parasite population originally isolated from Puerto Rico ([12,32], where parasites with high cercarial shedding show low virulence to the snail host. This inverse relationship between cercarial shedding and virulence was initially observed by Davies and colleagues [32] who isolated five inbred parasites lines from the SmPR1 laboratory line. Consistent with this, experimental selection of SmPR1 populations for high or low cercarial output resulted in rapid divergence in cercarial production, with high shedding parasites showing reduced virulence to the snail host. These results run counter to classical theoretical work suggesting that virulence is expected to be a byproduct of increased transmission stage production ([5,25]. However, these authors also showed that high virulence to the snail host was associated with lower virulence to the mammal host. They suggested an alternative trade-off model involving pleiotropy between genes underlying parasite traits conferring fitness within the definitive (mammal) and intermediate (snail) host [12]. They suggested that such pleiotropy might explain patterns of virulence observed and promote the maintenance of genetic and phenotypic polymorphisms in parasite populations utilizing multiple hosts. The intriguing differences in transmission stage production and virulence toward the snail host observed in work with SmPR1 [12,32] and our work suggest that the underlying causes of high virulence in different schistosome populations may vary [33].

We also demonstrated that the gender of the parasite can impact cercarial shedding. Indeed, for the SmBRE (LS) population only, we highlighted that male genotypes of *S. mansoni* produced significantly less cercariae than the female ones. This result is consistent with observations collected by Boissier and colleagues in their meta-analysis [34] of Brazilian schistosome/snail systems.

### What causes virulence to snails?

We compared virulence of our two schistosome populations by directly measuring mortality of inbred snails, and by quantifying laccase-like activity and hemoglobin rate. Schistosome parasites have a direct impact on their snail intermediate hosts: as they grow and generate cercariae, sporocysts deplete galactogen in the albumen gland and consume the ovotestis and hepatopancreas, converting stored glycogen to glucose [10,35]. Wang and colleagues [36] found that neuropeptides and precursor proteins involved in snail reproduction were heavily down regulated in infected prepatent snails compared to uninfected snails, suggesting that this could play a role in castration of *Biomphalaria* snails by schistosomes. Other down regulated neuropeptides in prepatent snails were linked to snail feeding and growth, process that directly impact the reproductive capacity, metabolism and immunity of the snail host. The high level of mortality observed in snails infected with SmLE (HS) is consistent with the fact that sporocysts cells comprised on average 26% to 47% of the cells within infected snails (cohort 2), largely replacing the hepatopancreas and ovotestis. This can also explain the reduction in laccase-like activity and hemoglobin rate in SmLE (HS) relative to SmBRE (LS), and the strong correlations between cercarial production, laccase-like activity and hemoglobin rate observed in SmLE (HS), as this rapidly growing parasite depletes snail host resources. Cercarial shedding is also harmful to snails because cercariae puncture the tegument to exit the snail, causing hemolymph loss, another potential cause of early death in infected snails (personal observations). This may also contribute to the higher virulence of SmLE (HS) compared with SmBRE (LS) population of parasites.

### Natural variation or an artifact of laboratory maintenance?

Our two populations of parasites (SmLE (HS) and SmBRE (LS)) have been maintained in laboratory conditions for 54 and 44 years respectively. We were concerned that long term maintenance of these parasites in the laboratory could have selected for the life history traits observed: for example serial passage of microbial pathogens often imposes selection for rapid growth and high virulence ([37–39]. Interestingly, cercarial production in SmLE and SmBRE has remained stable over multiple decades. Our SmLE (HS) population was reported to show high cercarial production compared to the other populations of *S. mansoni* parasites in a study published 33 years ago [11]. Similarly, the low shedding profile exhibited by our SmBRE (LS) population is consistent with low shedding by this *S. mansoni* population reported 26 years ago [15]. Hence, cercarial shedding phenotypes observed in these two parasite populations have remained stable over time.

### Dynamics of sporocyst growth

We observed dramatic differences in sporocyst growth profiles between SmLE (HS) and SmBRE (LS) using our qPCR assay (Figure 3A). These differences reflect sporocyst growth rate, rather than differences in number of infecting miracidia, because we exposed each snail to a single miracidium. The growth kinetics differs significantly: SmLE (HS) sporocysts have parabolic growth profile and a much higher proportion of daughter sporocysts produced, even during the prepatent period (i.e. 3 weeks after parasite exposure and 1 week before the first cercarial shedding). The SmLE (HS) *S. mansoni* population grows faster (reaching an average of 47% of cells within infected snails) compared to SmBRE (LS) population (reaching an average of 25% of cells within infected snails). Interestingly, the SmBRE (LS) daughter sporocyst kinetics in whole snails measured by qPCR is similar to that obtained by microscopy and 3D reconstructions of sporocysts (in the hepatopancreas only) in the same parasite population [15], with an initial exponential growth of the parasite tissues followed by a plateau.

Differences in sporocyst proportions in SmLE (HS) and SmBRE (LS) infected snails do not fully explain the difference in cercarial production between these lines. The SmBRE (LS) infected snails shed significantly fewer cercariae than predicted from qPCR measures of sporocysts cells in infected snails (Figure 3C). We suspect that SmLE (HS) and SmBRE (LS) sporocysts may exhibit different cellular trajectories, with differences in development of cell populations that differentiate to generate cercariae and those that give birth to the next generation of daughter sporocysts [40]. Advances in our understanding of stem cell differentiation of *S. mansoni* within the molluscan host now provide the tools needed to investigate these transmission related developmental differences at the cellular and molecular levels [40– 42].

## CONCLUSION

In this study, we describe two different transmission strategies of *S. mansoni* that have a strong genetic basis: a “boom-bust” strategy characterized by high virulence, high transmission and short duration infections for SmLE (HS), compared with “slow and steady” strategy with low virulence, low transmission but long duration of infection for SmBRE (LS) populations. We speculate that these two different strategies may be selected in the field to optimize the parasite transmission and fitness in different environments. We envisage that in high transmission areas, where individual snails may contain competing *S. mansoni* infections (or coinfections with other trematodes species [43], the SmLE (HS) strategy may be strongly selected as a consequence of intense within host competition. Conversely, in low prevalence sites, where coinfections are rare, the SmBRE (LS) strategy and limited virulence to the snail host may be advantageous. Genetic crosses between parasites from these two distinctive *S. mansoni* populations, followed by a classical quantitative trait locus analysis [44], now provides the opportunity to determine the genetic basis of these key transmission-related phenotypes in an important human helminth infection.

## DECLARATION

### Consent for publication

Not applicable

### Availability of data and materials

The datasets generated and/or analyzed during the current study are available from the Zenodo repository, [PERSISTENT WEB LINK TO DATASETS]

### Authors’ contributions

WL, FDC and TJCA designed the experiments. WL, RD, FDC and MMW performed the experiments. WL performed the data analyses. WL and TJCA drafted the manuscript. All authors read and approved the final manuscript.

## Acknowledgements

We thank Michael S. Blouin from Oregon State University for providing the Bg26 snail line, Guillaume Mitta and Benjamin Gourbal (University of Perpignan France) for providing SmBRE *S. mansoni* population and Philip LoVerde from UT Health (San Antonio) for providing SmLE *S. mansoni* population.

## FIGURE LEGENDS

**Supplementary figure 1: Multiplex PCR assay for identifying infected prepatent snails.** We electrophoresed multiplexed PCR products generated using *piwi* and *α-tubulin-2* primers on 2% agarose gel. The size ladder used is the 100 bp ladder from Promega. Infected *B. glabrata* Bg26 snails show a “double-band”: a 361 bp *piwi* snail specific band and a 190 bp *α-tubulin-2* parasite specific band. Uninfected snails exhibit only the 361 bp *piwi* snail specific band while *S. mansoni* control show only the 190 bp *α-tubulin-2* parasite specific band.

**Supplementary figure 2: Impact of *S. mansoni* gender on the cercarial production.** Male sporocysts produced significantly less cercariae than female sporocysts in SmBRE (LS) parasite. There were no difference driven by the gender of the parasites for the SmLE (HS) population of *S. mansoni*. \**p* < 0.05; ** *p* ≤ 0.01; *** *p* ≤ 0.001.

**Supplementary figure 3: Virulence of *S. mansoni* parasites: correlation between cercarial production and measured *B. glabrata* snail physiological parameters. (A)** There is a negative correlation between the average of cercariae produced by a snail and the total laccase-like activity in the hemolymph of this snail (Pearson’s test, coef. = −0.33). **(B)** Similarly, the hemoglobin rate is negatively correlated to the cercarial output (Pearson’s test, coef. = −0.54). **(C)** We also observed a strong positive correlation between the total laccase-like activity and the hemoglobin rate in the hemolymph of the snails. Both of these parameters are good proxies for assessment of snail health (Pearson’s test, coef. = 0.78).

