## Supplementary figures and images for "Striking differences in virulence, transmission, and sporocyst growth dynamics between two schistosome populations"

### Supplementary figure 1

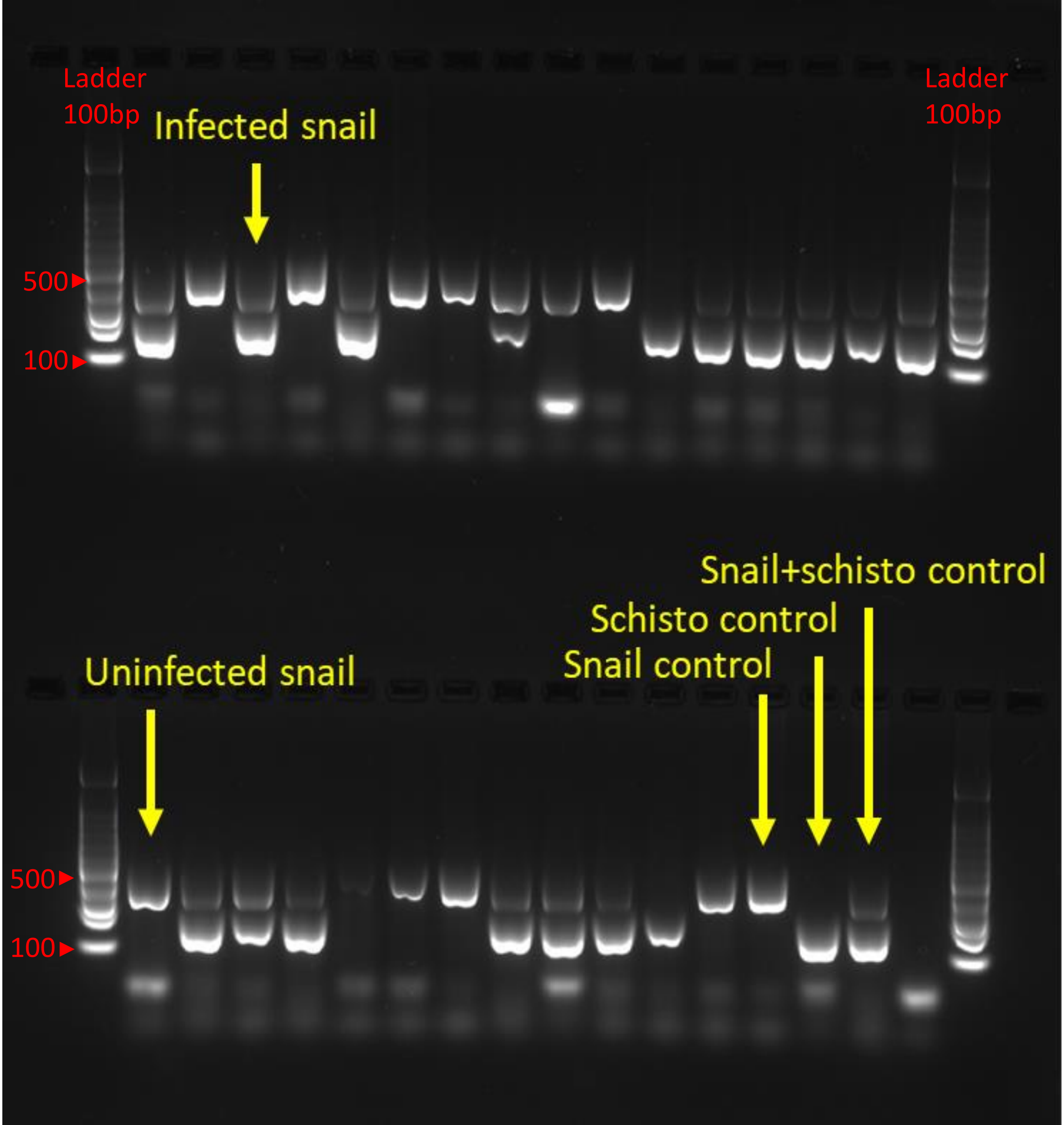

### Supplementary figure 2

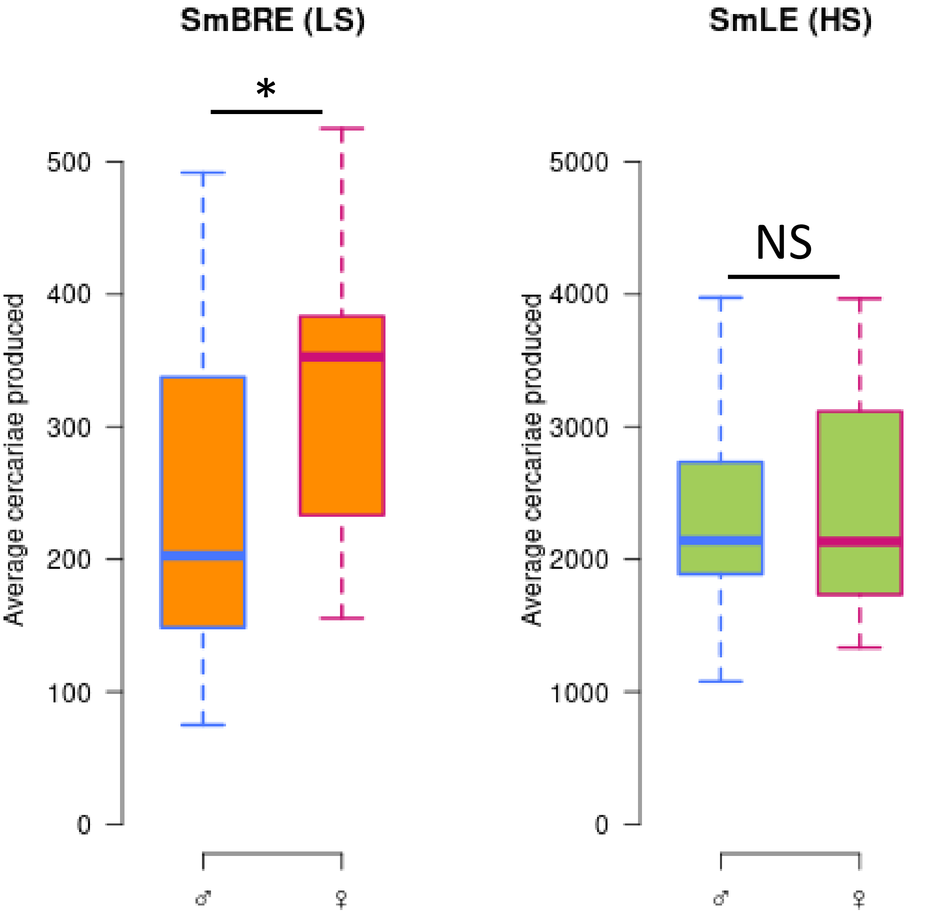

### Supplementary figure 3

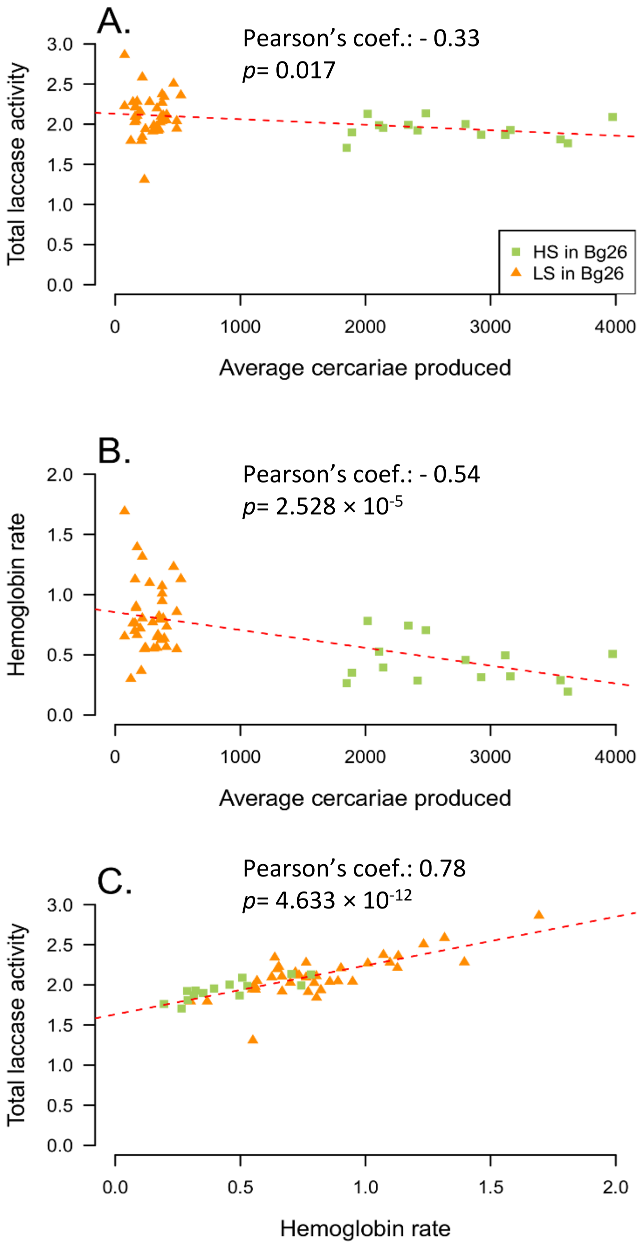
